# Evolution of Hominin Polyunsaturated Fatty Acid Metabolism: From Africa to the New World

**DOI:** 10.1101/175067

**Authors:** Daniel N. Harris, Ingo Ruczinski, Lisa R. Yanek, Lewis C. Becker, Diane M. Becker, Heinner Guio, Tao Cui, Floyd H. Chilton, Rasika A. Mathias, Timothy D. O’Connor

**Affiliations:** Institute for Genome Sciences, University of Maryland School of Medicine, Baltimore, MD; Department of Medicine, University of Maryland School of Medicine, Baltimore, MD; Program in Personalized and Genomic Medicine, University of Maryland School of Medicine, Baltimore, MD; Department of Biostatistics, Johns Hopkins Bloomberg School of Public Health, Baltimore MD; GeneSTAR Research Program, Johns Hopkins University School of Medicine. Baltimore, MD 21287; Laboratorio de Biología Molecular, Instituto Nacional de Salud, Lima, Perú; Department of Urology, Wake Forest School of Medicine, Winston-Salem, NC; Department of Physiology/Pharmacology, Wake Forest School of Medicine, Winston-Salem, NC; Department of Cancer Biology, Wake Forest School of Medicine, Winston-Salem, NC

**Keywords:** Evolution, Ancient DNA, Population Genetics, Polyunsaturated Fatty Acids

## Abstract

**Background:** The metabolic conversion of dietary omega-3 and omega-6 18 carbon (18C) to long chain (> 20 carbon) polyunsaturated fatty acids (LC-PUFAs) is vital for human life. Fatty acid desaturase (*FADS*) 1 and 2 catalyze the rate-limiting steps in the biosynthesis of LC-PUFAs. The *FADS* region contains two haplotypes; ancestral and derived, where the derived haplotypes are associated with more efficient LC-PUFA biosynthesis and is nearly fixed in Africa. In addition, Native American populations appear to be nearly fixed for the lesser efficient ancestral haplotype, which could be a public health problem due to associated low LC-PUFA levels, while Eurasia is polymorphic. This haplotype frequency distribution is suggestive of archaic re-introduction of the ancestral haplotype to non-African populations or ancient polymorphism with differential selection patterns across the globe. Therefore, we tested the *FADS* region for archaic introgression or ancient polymorphism. We specifically addressed the genetic architecture of the *FADS* region in Native American populations to better understand this potential public health impact.

**Results:** We confirmed Native American ancestry is nearly fixed for the ancestral haplotype and is under positive selection. The ancestral haplotype frequency is also correlated to Siberian populations’ geographic location further suggesting the ancestral haplotype’ s role in cold weather adaptation and leading to the high haplotype frequency within Native American populations’. We also find that the Neanderthal is more closely related to the derived haplotypes while the Denisovan clusters closer to the ancestral haplotypes. In addition, the derived haplotypes have a time to the most recent common ancestor of 688,474 years ago which is within the range of the modern-archaic hominin divergence.

**Conclusions:** These results support an ancient polymorphism forming in the *FADS* gene region with differential selection pressures acting on the derived and ancestral haplotypes due to the old age of the derived haplotypes and the ancestral haplotype being under positive selection in Native American ancestry populations. Further, the near fixation of the less efficient ancestral haplotype in Native American ancestry suggests the need for future studies to explore the potential health risk of associated low LC-PUFA levels in Native American ancestry populations.

## Background

The metabolic conversion of omega-3 (n-3) and omega-6 (n-6) dietary 18 carbon (18C) polyunsaturated fatty acids (PUFAs) to biologically active long-chain PUFAs (> 20 carbon, LC-PUFAs) is essential for human life. LC-PUFAs and their metabolites are vital structural and signaling components for numerous biological systems including brain development and function, innate immunity and energy homeostasis [1, 2]. Consequently, the capacity of populations to adapt to their PUFA environments and synthesize or ingest LC-PUFAs is essential to their survival [3-6].

In the modern Western diet (MWD), the majority (> 90%) of all PUFAs consumed are two typically plant sourced 18C-PUFAs, α–linolenic acid (ALA, 18:3n-3) and linoleic acid (LA, 18:2n-6). Over the past 50 years, the ingestion of LA dramatically increased (∼3 fold, 6-8% of daily energy consumed) due to the addition of vegetable oil products (e.g. soybean, corn, and canola oils; and margarine/shortenings) to the MWD [7]. Once ingested, n-6 and n-3 18C-PUFAs can be converted into several LC-PUFAs including eicosapentaenoic acid (EPA, 20:5n-3), docosapentaenoic acid (DPA, 22:5n-3), docosahexaenoic acid (DHA, 22:6n-3) and arachidonic acid (ARA, 20:4n-6) utilizing desaturase and elongase enzymes [3]. There are also dietary sources of preformed LC-PUFAs with eggs and certain meats containing ARA; and seafood being highly enriched for DHA, EPA and DPA [7-9].

The *FADS* gene region (chr11: 61,540,615-61,664,170) contains *FADS1*, and *FADS2* which encode for desaturase enzymes that catalyze the rate-limiting steps in converting 18C-PUFAs into LC-PUFAs [10]. It was originally assumed that LC-PUFAs biosynthesis from 18C-PUFAs was highly inefficient and similar in all human populations [11]. However, numerous studies revealed that levels of LC-PUFAs and metabolic efficiencies by which LC-PUFAs are formed are highly impacted by an individual’ s genetic background [12-16].

Ameur et al. [16] found two regions of high LD in Europeans, with the first block spanning *FADS1* and part of *FADS2*. This region has two major haplotypes, with the derived haplotype associated with more efficient conversion of 18C-PUFAs into LC-PUFAs and being most common in African populations. Native American populations appear to be fixed for the ancestral haplotype that is associated with lower *FADS* activities and levels of LC-PUFAs [15, 16]. However prior analyses only had access to small Native American ancestry sample size [15, 16] which did not facilitate a comprehensive examination of the genomic architecture of the *FADS* gene region in Native American ancestry populations. In addition, more recent studies suggest that the ancestral haplotype is under positive selection in the Greenlandic Inuit [17] and Native American populations [18], while the derived haplotype is under positive selection in African [15, 16] and European populations [19]. These findings provide evidence that differential selection pressures that favor one *FADS* haplotype over another may play a critical role in the capacity of populations to adapt to regional environments. Additionally, the fact that African populations are nearly fixed for the derived haplotype while Eurasian populations are polymorphic [13, 15, 16] and the derived haplotype is largely absent in Native American populations creates an interesting evolutionary puzzle, which could be explained by archaic reintroduction of the ancestral haplotype in non-African populations [20].

Modern humans mated with archaic hominins such as Neanderthals and Denisovans, after migrating out of Africa. Specifically this is supported by the presence of archaic haplotypes in non-African genomes [21-26]. Some of these introgressed haplotypes are associated with modern human phenotypes and diseases [27-29], giving emphasis to how this evolutionary history impacted modern humans. In addition, Neanderthals and Denisovans appear to be homozygous for the *FADS* gene cluster ancestral haplotype [16]. Since African populations are nearly fixed for the derived haplotype while non-African populations are polymorphic [13, 15, 16], it is feasible that the derived haplotype rose to fixation in humans prior to “out-of-Africa”, and then the ancestral haplotype was re-introduced to non-Africans through admixture with archaic hominins in Eurasia.

An alternative hypothesis is that the derived haplotype began to form prior to the divergence of Neanderthals, Denisovans, and modern humans which was followed by differential selection pressures in different environments. There is great genetic difference between the human derived and ancestral haplotypes [15, 16] which suggests these haplotypes are old in the human lineage. If the derived haplotype began to form near the divergence of these three hominins, then there would be at least 550,000 years [22] for more mutations to occur between the two haplotypes. Previous estimates for the time to the most recent common ancestor (TMRCA) of all *FADS* human haplotypes is 1.49 million years ago (ya) [15] and TMRCA of the derived haplotype is as old as 433,000 ya [16]. However a recent analysis examining the archaic haplotype topology suggest differential relations of the Neanderthal and Denisovan haplotypes with the modern human ancestral and derived haplotypes [19]. Therefore, it is plausible that the TMRCA of the derived haplotype is in fact older than the modern-archaic human divergence. Further, the differential selection pressures of the ancestral and derived haplotypes [15, 17-19] could then explain the global haplotype frequency distribution.

Therefore we analyzed this genetic region for signs of archaic admixture or ancient development of the derived haplotype through the use of the 1000 Genomes Project [30], European American and African American genomes from GeneSTAR [13], as well as Native American ancestry individuals from the Peruvian Genome Project (Unpublished Data^a^). Further, due to the importance of LC-PUFAs to human health, we placed an added focus on Native American ancestry individuals to further illuminate the genomic architecture of this region in Native American ancestry populations.

## Methods

### 1. Data Preparation

We jointly called all positions in the range chr11:61,540,615-61,664,170 genome build hg19 [30] in 127 African American and 156 European American genomes from GeneSTAR[13], 67 Native American and 47 mestizo (Native American-European admixed ancestry) genomes from the Peruvian Genome Project (Unpublished Data^a^) and the Neanderthal [22] and Denisovan [21] genome alignments downloaded from the Max Planck Institute for Evolutionary Anthropology, using GATK UnifiedGenotyper [31]. We removed all invariant sites, variant calls flagged as LowQual, sites given a quality score of ≤ 20, INDELS, or triallelic single nucleotide polymorphisms (SNPs) with vcftools v0.1.11 and PLINK1.9 [32-34] which returned a high coverage genome dataset with 2,122 high quality bialleic SNPs. We then intersected the same genomic region variant calls from all 2,504 individuals from the 1000 Genomes Project [30] with the high coverage dataset, to create a more diverse but low coverage dataset. We removed triallelic SNPs that were not created by a strand flip and flipped the strand in the 1000 Genomes Project genomes to correct those that were created by a strand flip, with PLINK1.9 [32, 33]. After the merge with 1000 Genomes Project and filters, 899 variants remained. Some analyses used the high coverage dataset while others used the combined low coverage diverse dataset. The following methods sections will indicate which dataset was used for each analysis by referring to the dataset used as high coverage or low coverage.

### 2. Phasing and Local Ancestry Inference

The genomes in the low and high coverage dataset were phased with SHAPEIT v2.r790 [35] separately, using default settings. All missing genotype positions were imputed by SHAPEIT and for all analyses except for local ancestry, were set back to missing. Local ancestry was calculated in the low coverage dataset on Native American and mestizo genomes with < 99% Native American ancestry (Unpublished Data^a^), the admixed populations from the 1000 Genomes Project [30], and all individuals from GeneSTAR [13] using the default setting of RFMix [36]. We used Native American and mestizo individuals with ≥ 99% Native American ancestry (Unpublished Data^a^****) as the Native American reference, and the CEU and YRI as the European and African reference respectively [30].

### 3. GeneSTAR Relatedness Filtering

All data sources except GeneSTAR were already filtered for kinship to remove at least 3^rd^ degree relations [30] (Unpublished Data^a^). Pedigree information was used to remove individuals so that no closer than 3^rd^ degree relations remained in the African American and European American families in GeneSTAR [13]. All the pedigree information was previously validated with genome wide array data and genetic kinship analysis [37]. This kinship filtering resulted in 101 African American and 128 European American individuals to yield a total of 229 unrelated GeneSTAR samples. All analyses detailed bellow that include GeneSTAR samples only include the 229 unrelated samples.

### 4. Ancestral and Derived Haplotype Proportion Calculations

Haplotypes were determined as either ancestral or derived from the genotype at SNP rs174537 (chr11:61,552,680) which is the most representative of the haplotype within Africa [15]. Haplotype proportions were calculated in all populations from the low coverage dataset. In addition, we binned the haplotype proportions by the estimated local ancestry haplotypes. This was done on these individuals with (380 African, 450 European, and 216 Native American estimated haplotypes) and without (505 African, 685 European, and 386 Native American estimated haplotypes) homozygous local ancestry calls.

### 5. Analysis for Signs of Selection on the *FADS* Haplotype in Peruvians

We calculated the population branch statistic (PBS) to determine if the ancestral haplotype is under selection in Native Americans relative to the CEU and the CHB [38]. An additional dataset was first created by taking the Peruvian Genome Project autosome genotype calls and merging with the 1000 Genome Project using the same merging and filtering procedure as mentioned above. To calculate the PBS we used PLINK1.9 to calculate F_ST_ between Native Americans from the Peruvian Genome Project (NatAm) (Unpublished Data^a^), CEU, and CHB from the 1000 Genomes Project [30] over the entire genome [32, 33]. We then computed the PBS statistic of the form:

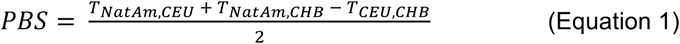

throughout the entire autosome, where *T*_*NatAm,CEU*_ *, T*_*NatAm,CHB*_ and *T*_*CHU,CHB*_ represent the F_ST_ log-transformed time value calculated between the Native American and CEU, Native American and CHB, and CEU and CHB populations respectively. A Z-score was computed for all SNPs within +/-500 kb of the *FADS* gene region by comparing to the genome wide average and its standard deviation. The Z-score was then converted to a P-value with a correction for multiple hypothesis testing. We calculated all combinations of the PBS statistic such that SNPs were assessed for being under selection in the CEU and CHB, which were plotted as branch lengths.

### 6. Siberian Genome Analysis

Genotype data from 12 Siberian populations [39] were merged with the Human Genome Diversity Panel (HGDP) [40] and we removed all tri-allelic sites with plink [32, 33]. The ancestral and derived haplotype proportion was calculated using rs174537. We used ADMIXTURE [41] to calculate global ancestry proportions in all Siberian, East and South Asian populations. In addition, we included the Yoruba and French populations to serve as the African and European ancestry reference respectively. The autosome was first filtered by removing singletons and sites with > 10% missing genotypes. We then LD pruned the data with PLINK1.9 indep-pairwise 50 5 0.5 [32, 33]. ADMIXTURE [41] was run on K 1-10, randomly 10 times for each K. K=6 was selected as the most representative of these data due to K=6 having the lowest cross validation. We correlated Siberian, and East and South Asian ancestral haplotype proportion to their latitude coordinates independent of their European admixture, as determined by the ADMIXTURE estimates, by computing the linear regression: lm(proportion ∼ European_admixture + latitude) and only analyzed the P value relative to the correlation with latitude. Correlation to temperature would be a better comparison, however temperature data is unavailable for these populations and many of their geographic locations are remote. Therefore we cannot use a nearby city to obtain temperature data for these populations. However, the general trend from South Asia to Siberia is a decrease in average yearly temperature [42] which supports using latitude as an appropriate indication of temperature for this comparison.

### 7. Ancient Humans

We downloaded SRA files from the Mota [43], ancient Eskimo [39], Anzick-1 [44], MA-1 [45] genomes and extracted fastq files with sra-toolkit fastq-dump [46] from the Eskimo, Anczck-1, and MA-1 genomes and used sam-dump [46] to extract a sam file for the Mota individual. We downloaded bam files for all ancient European genomes [44, 47-50] and the Ust’ Ishim individual [51]. Single read and paired-end read fastq files were generated for the Ust’ Ishim individual and single read fastq files were generated for the Stuttgart and Loschbour ancient Europeans using Bedtools version 2.17.0 BAMtoFASTQ [52]. All fastq files were aligned to hg19 [30] with bwa mem [53]. The paired-end and single reads were aligned to hg19 separately with bwa mem and then combined into one bam file using samtools v0.1.19-44428 merge [54] for the Ust-Ishim individual. Duplicates were marked in the Stuttgart and Loschbour bam files with Picard Tools version 1.79 MarkDuplicates [55]. The Mota individual sam file was re-formatted to be compatible with GATK [31] and was then converted to a bam file with samtools view. All bam files were sorted and indexed with samtools [54]. We called all positions in the *FADS* gene region in each ancient genome individually with GATK UnifiedGenotyper [31]. Each individual was genotyped as homozygous ancestral or derived, or heterozygous based on the genotype of rs174537.

### 8. Recombination Mapping

To determine the effectiveness of rs174537 tagging the ancestral and derived haplotypes, we used the R package rehh bifurcation.diagram function [56] to perform recombination mapping in the low coverage dataset. rs174537 was set as the variant to examine how the haplotype decayed within the *FADS* region due to recombination on the ancestral or derived haplotypes.

### 9. Haplotype Construction and Network Analysis

We used PLINK to form LD blocks in the Native American samples with PLINK1.9’ s hap command [32, 33]. We then selected the region chr11:61,543,499-61,591,907 with the high coverage dataset or chr11:61,543,499-61,591,636 with the low coverage dataset to construct haplotype networks. We constructed haplotype networks using the R package pegas [57]. The human-chimpanzee ancestral reconstructed reference sequence represented the outgroup haplotype [30]. We loaded all DNA sequences into R using read.dna from the R package ape [58], then formed the haplotypes using haplotype and a network using haploNet, both from the R package pegas [57]. We plotted the haplotype network using the normal R plot function. Using the described haplotype network method, we constructed haplotype networks removing all haplotypes with a count of ≤ 3 (except for the Denisovan and Neanderthal haplotypes and the human-chimpanzee ancestral reconstructed reference haplotype). Haplotype networks were then colored by ancestral vs derived, or local ancestry (if calculations available) and global population ancestry (where local ancestry calculations were unavailable). The high coverage haplotype network also included modern human invariant sites so that sites variable in the archaic hominins and the human-chimpanzee ancestor relative to modern humans could be compared.

### 10. Tree Analysis

To convert the haplotype network into a tree we calculated all pairwise differences between each haplotype to form a matrix of differences between all haplotypes in the high coverage dataset. Then Phylip neighbor v 3.68 was used to form a neighbor-joining tree based on the matrix of differences between each haplotype [59]. To assign confidence that each node is correct we performed 500 bootstraps by randomly sampling, with replacement, each base over the entire haplotype length for each haplotype. Phylip consense [59] was used to calculate a consensus tree from the 500 bootstraps with the human-chimpanzee ancestral reconstructed reference sequence set as the root of the tree [59]. We then used MEGA7 to plot the consensus tree [60].

### 11. Derived Haplotype TMRCA Calculations

To determine when the derived haplotype arose, we calculated the TMRCA based on the degree of differentiation from a human ancestor sequence described in the following equation:

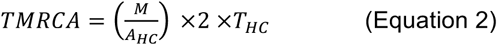

from Coop et al. [61], where *M* is the average number of differences between all human derived haplotypes and the human-chimpanzee ancestral reconstructed reference sequence [30] over the entire high quality ancestral call bases of the haplotype length (47,820 bases out of the total haplotype length of 48,408 bases). Only mutations that follow the infinite sites model [62] in modern humans were used for this analysis, meaning only mutations found in the derived haplotypes and not present in the ancestral haplotypes were used. *A*_*HC*_ represents the local human chimpanzee divergence value which we calculated, to be 0.829%, by using liftover [63] to determine the hg19 *FADS* haplotype coordinates in the panTro-4 reference genome downloaded from the UCSC Genome Browser [64, 65]. We then aligned the chimpanzee and human reference sequences for the two regions with Clustal Omega 2.1 [66] and calculated the percentage of mutations between the human and chimpanzee. The value *T*_*HC*_ corresponds to the time of the human-chimpanzee divergence, which we specified as 6,500,000 ya [67-69]. We then calculated the variance and standard deviation of the TMRCA for all derived haplotypes and applied the framework by Hudson [70] to calculate the 95% confidence interval (CI). This analysis only used the high coverage dataset which included modern human invariant positions. We assessed the impact of differing human-chimpanzee divergence values by keeping *M* and *A*_*HC*_ constant while varying *T*_*HC*_ between 5,000,000 to 7,000,000 ya.

## Results

### 1. Organization of the *FADS* Gene Region

Recombination mapping shows that there is one major haplotype for both the ancestral and derived haplotype over the entire *FADS* gene region based on rs174537 (Additional File 1: Figure S1). However, prior research shows that there are two sub-regions within the *FADS* region and that the region chr11:61,567,753-61,606,683 is associated with increased biosynthesis of LC-PUFAs [16]. We re-define this region to be chr11:61,543,499-61,591,907 through plink LD block formation using the Peruvian Genome project samples to isolate the largest LD block. With the high coverage GeneSTAR and Peruvian Genome Project samples, this region contains 43 SNPs. When intersecting with the 1000 Genomes samples, this region only contains 38 SNPs (Additional File 2: Table S1) in the region: chr11:61,543,499-61,591,636.

### 2. Ancestral and Derived Haplotype Proportion

Global haplotype proportions confirm that African populations are nearly fixed for the derived haplotype and Eurasia is polymorphic [13, 15, 16] (Figure 1A). The GeneSTAR African and European Americans have similar haplotype proportions as populations in the 1000 Genomes Project from those same regions. Siberian and the Peruvian Genome Project populations have the lowest average derived haplotype proportions, which is consistent with the hypothesis that the ancestral haplotype confers cold weather adaptation [17]. The 1000 Genomes American populations have a higher ancestral haplotype frequency than Eurasian populations. However, with 59 of the 114 individuals from the Peruvian Genome Project being ≥ 99% Native American ancestry (Unpublished Data^a^), we are more appropriately positioned to address the genetic architecture of this region in Native Americans. Ancestral haplotype proportions are greater in the mestizo Peruvians, than any 1000 Genomes populations. Native American identifying populations have an even higher ancestral haplotype proportion than the mestizo Peruvians (Figure 1A). However neither Peruvian population is fixed for the ancestral haplotype. This is likely due to European admixture in these samples where they contain the derived haplotype within a European haplotype. To examine this phenomenon, we calculated local ancestry in all admixed 1000 Genomes African American and mestizo populations in addition to GeneSTAR and Peruvians with < 99% Native American ancestry. This shows that the ancestral haplotype is nearly fixed in Native American ancestry as 97.44% of 386 haplotypes have the ancestral haplotype (Figure 1B). Further, derived Native American haplotypes come from 1 African American, 3 European Americans, 1 Mexican, and 4 Puerto Rican individuals. Only one of these individuals, a Puerto Rican individual, was homozygous for Native American ancestry. This indicates that recombination could have disrupted the Native American haplotype in the Mexican and three Puerto Rican individuals that were heterozygous in local ancestry calls to introduce the derived haplotype because the other haplotype was European. The same could have occurred in the African American and European American individuals, however it is also likely that this is an error in local ancestry inference since these populations are expected to have little to no Native American ancestry [30]. Interestingly there is one Puerto Rican individual who is homozygous for Native American ancestry but is heterozygous at rs174537. This indicates that Native American ancestry is not completely fixed for the ancestral haplotype. When we restrict our calculations to those individuals with unambiguous ancestry (ie homozygous for a single ancestry) we find Africa is 99.74% for the derived and Native American is 99.54% for the ancestral haplotype (Figure S2) which further shows that both ancestries are nearly fixed for the derived or ancestral haplotype. European local ancestry haplotypes’ derived haplotype proportions follow the observed European population haplotype proportion patterns (Figure 1, Additional File 3: Figure S2).

**Figure 1.**
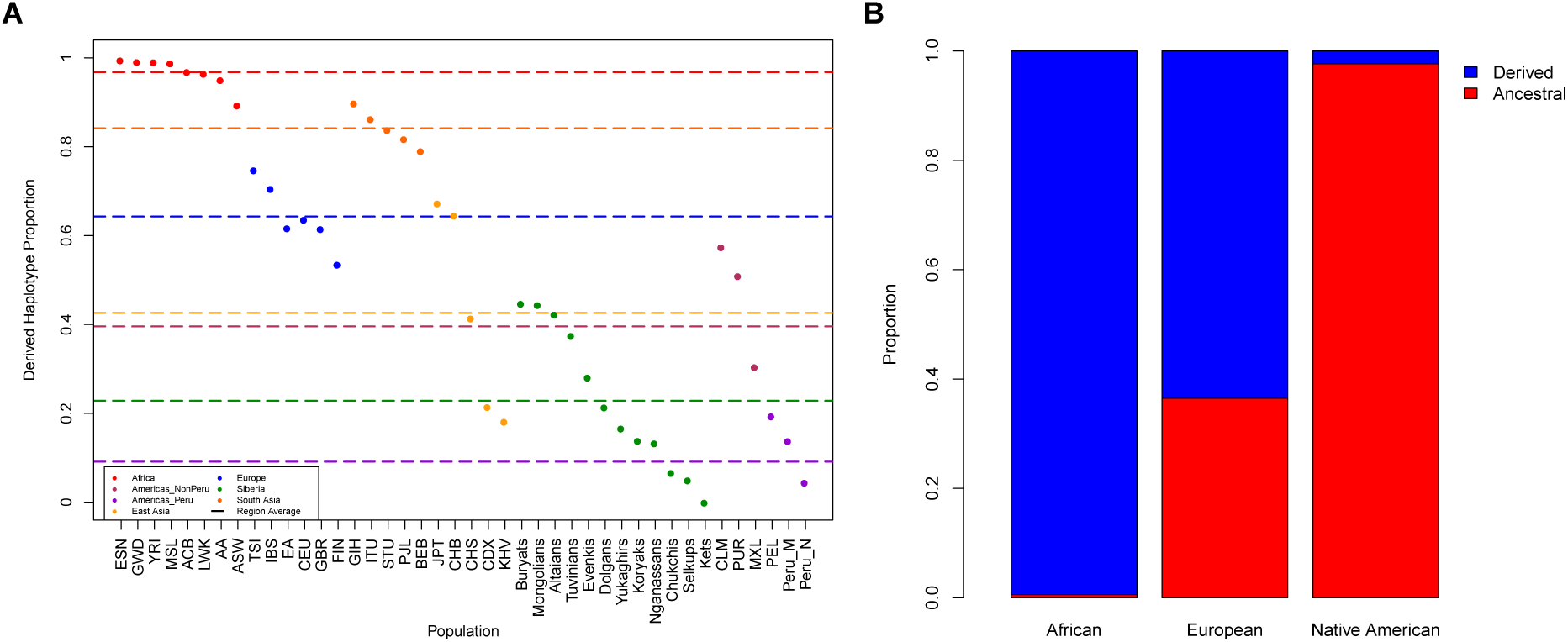
*FADS* ancestral and derived haplotype proportions in modern human populations. A) Derived haplotype proportion from all 1000 Genomes, Siberian, GeneSTAR African American (AA) and European American (EA) and Peruvian mestizo (Peru_M) and Native American populations (Peru_N). Dashed lines represent the geographic region derived average proportion. Americas_NonPeru consists of the 1000 Genomes Native American ancestry populations and Americas_Peru represents the Peruvian Genome Project populations. B) Ancestral (red) and derived (blue) haplotype proportion in all local ancestry segments from the admixed 1000 Genomes Project populations, African and European Americans from GeneSTAR, and Peruvian Genome Project individuals with < 99% Native American ancestry. (African = 505, European = 685, Native American = 386 haplotypes).

### 3. Selection in Native Americans and Cold Weather Adaptation in Siberians

Fumagalli et al. [17] showed that the *FADS* region is under positive selection in the Greenlandic Inuit. Recently, Amorim et al. [18] demonstrated the *FADS* cluster is under positive selection in Native American populations. We sought to perform a replication analysis to determine if the *FADS* region shows signs of positive selection in Peruvians. rs174537, and the surrounding *FADS* region, shows evidence of being under positive selection for the ancestral haplotype in Native Americans relative to the CHB and CEU (PBS = 0.33, P = 0.0001242) (Figure 2). The results from Greenland [17], other Native American populations [18], and now Peru indicate that the ancestral haplotype was beneficial to ancestral populations that migrated to the Americas. Fumagalli et al. [17] suggested that the ancestral haplotype is beneficial for cold weather adaptation by assisting with responding to dietary restricitions imposed by such a climate. We tested this by exploring the correlation between Siberian populations and their geographic cooridinates and found a sigificant correlation independent of European admixture (β= 0.01016, R^2^ = 0.3929, P = 5.06 x 10^-5^), such that the ancestral haplotype is at a higher proportion in more Northern regions (Additional File 4: Figure S3, Additional File 5: S4). This supports the ancestral haplotype being involved in cold weather adaptation, potentially as a response to dietary restrictions [17]; however, future studies are required to determine a molecular mechanism.

**Figure 2.**
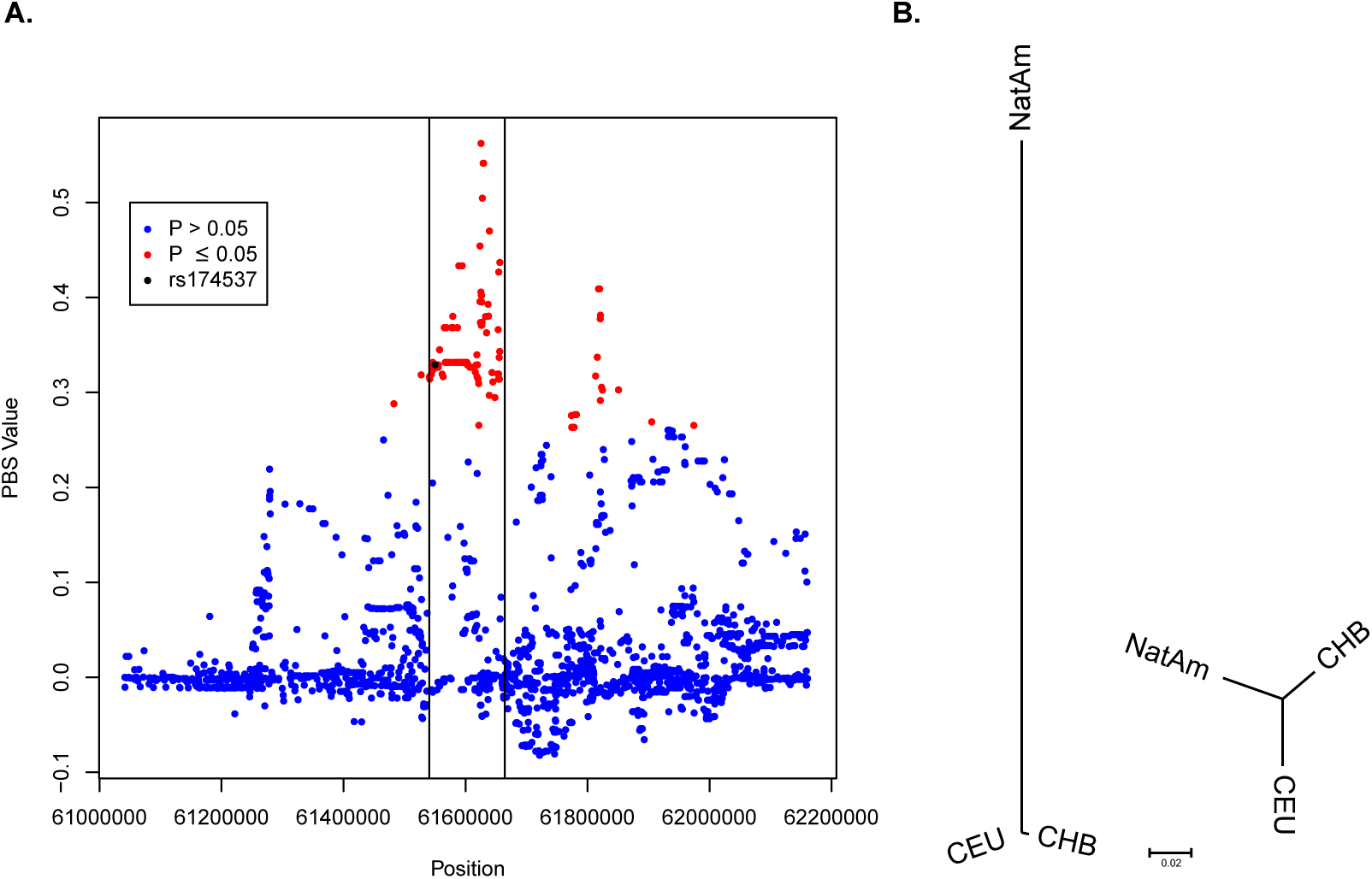
Signatures of selection in the *FADS* region in Native Americans. A) PBS values from chr11: 61,044,208-62,163,145. The black lines ma the *FADS* gene region and the black data point represents the PBS value for rs174537 (P = 0.0001242) while red points represent P values that a ≤ 0.05 after correction for multiple hypothesis testing. B) Representation of rs174537 PBS values (left) and the genome wide average PBS valu (right). Branch lengths are the PBS values for each population.

### 4. Ancient Humans

To better understand the evolution of this haplotype, we tested if ancient humans follow the pattern of haplotype proportions seen in modern humans. The ancient African Mota individual [43] was found to be homozygous for the derived haplotype (Additional File 6: Table S2) which is expected due to near fixation of the derived haplotype in African populations (Figure 1). Analysis of 19 Neolithic and Bronze age Europeans [48] revealed the haplotype to be polymorphic in ancient Europe (Additional File 6: Table S2). We analyzed two ancient individuals found in Siberia to understand how this haplotype travelled with Native American ancestors. A 45,000 year old Siberian [51] was found to be homozygous for the ancestral haplotype and even more significantly the Mal’ ta-1 individual [45] was also found to be homozygous for the ancestral haplotype (Additional File 6: Table S2). Mal’ Ta-1 is representative of a population that contribtuted to the Native American founding population’ s gene pool [45]. This indicates that the founders of the Americas also had this haplotype at a high proportion. In addition, an ancient Eskimo from Greenland whose ancestry was determined to be due to a later migration into the Americas from Siberia [39] was also homozygous for the ancestral haplotype (Additional File 6: Table S2). Further, the Anzick-1 individual from Montana [44] is homozygous for the ancestral haplotype (Additional File 6: Table S2). Since this individual is representative of the initial wave of migration into the Americas [44], it further indicates that the ancestral haplotype was likely at a high proportion in the founding population of the Americas. However, due to small sample size of ancient individuals, it is impossible to determine the haplotype proportion of the ancestral haplotype or if it was fixed or near fixed in the population that founded the Americas with ancient genomes.

### 5. *FADS* Haplotype Topography in Modern Humans and Archaic Hominins

Haplotype networks revealed two main clusters, a derived and ancestral cluster with the archaics intermediate (Figure 3). Further, the population local ancestry (Figure 1B) replicated the haplotype proportions distribution as the majority of Native American haplotypes appear in the ancestral cluster and the majority of African haplotypes in the derived with Eurasian haplotypes distributed among both clusters (Figure 3, Additional File 7: Figure S5). There is not one unique signature representing all Native American ancestral haplotypes, Eurasian, or African derived haplotypes (Table 1). A core haplotype for these three can be formed, however the core haplotypes contain variants found in other ancestral or derived haplotypes (Table 1). This supports the large degree of diversity of ancestries seen in the haplotype network (Additional File 7: Figure S5). For example, there are few Native American specific haplotypes; however, there are multiple Native American-Eurasian ancestral haplotypes (Figure 3, Additional File 7: Figure S5). This indicates that the ancestral haplotypes were present in the ancestral population of the Americas, and rose to high frequency either in Beringia or once in the new world.

**Figure 3.**
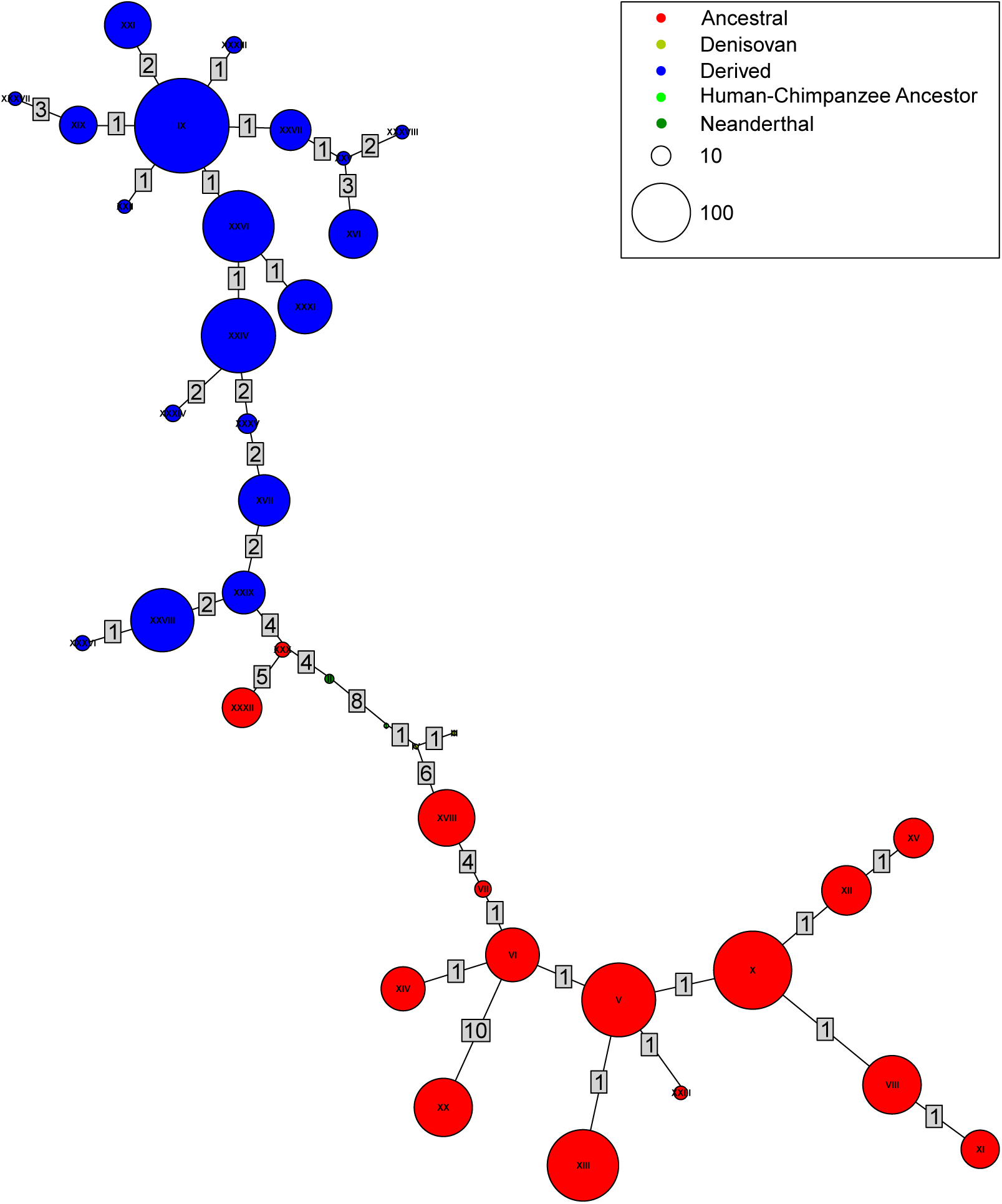
Haplotype network of chr11:61,543,499-61,591,636. All haplotypes are at least a frequency of 4 except for the Neanderthal, Denisovan, and Human-Chimpanzee Ancestor haplotypes. Numbers present on each link represent the number of mutations that separate the two haplotypes.

**Table 1.**
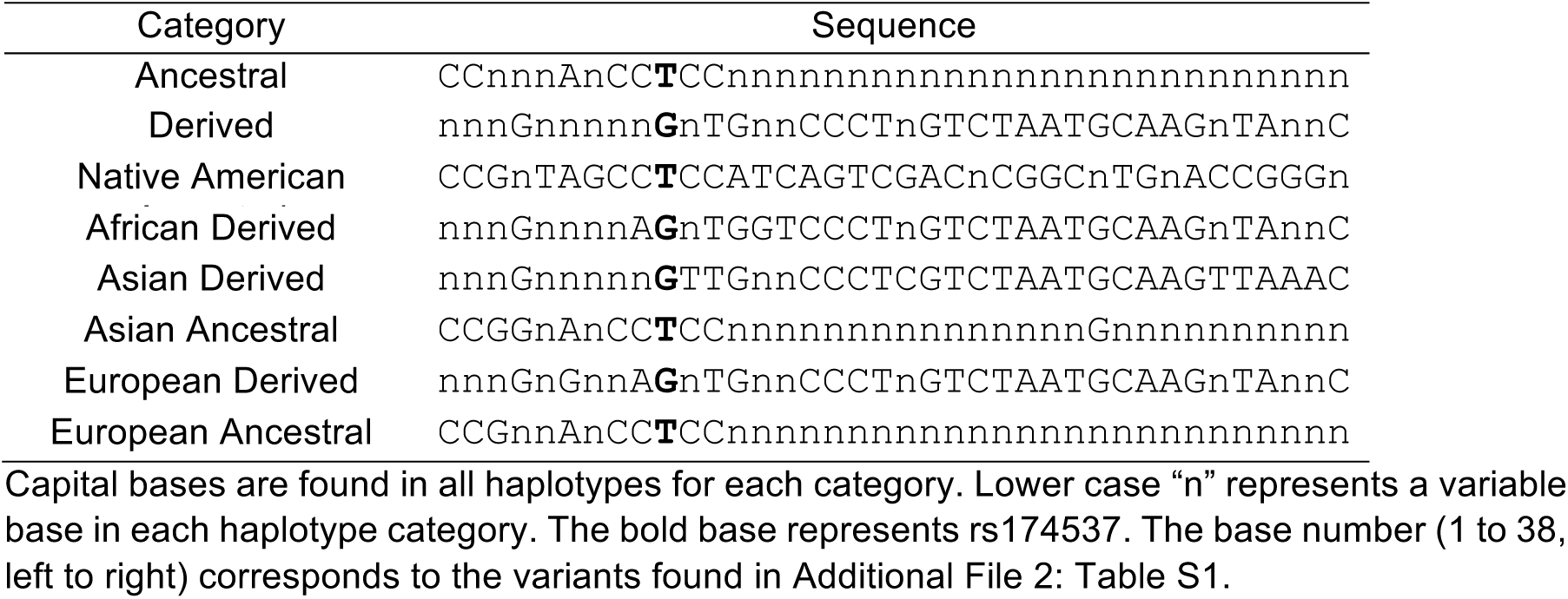
Core Haplotypes.

Interestingly, there are two ancestral haplotypes that are closer to the derived cluster than the ancestral cluster (HAP# XXX, XXXII) (Figure 3). Haplotype XXXII is likely due to recombination merging a derived and ancestral haplotype as all individuals have either Asian or European ancestry (Additional File 7: Figure S5). Haplotype XXX contains one Colombian individual who has African ancestry in the ancestral haplotype and European ancestry in their other haplotype (Figure 3, Additional File 7: Figure S5). Therefore this individual could also have recombination bringing together the derived and ancestral haplotype. The other individuals come from continental African populations in the 1000 Genomes project (Additional File 7: Figure S5), which have little European admixture [30], therefore making this unlikely to be recombination. Instead this could represent an evolutionary intermediate between the ancestral and derived haplotype, however functional studies of this haplotype are required to determine the efficiency of conversion of 18C-PUFAs to LC-PUFA metabolism.

Due to the global distribution of the derived and ancestral haplotype frequencies this locus is a candidate for archaic re-introduction of the ancestral haplotype to non-Africans. However these networks do not present a haplotype topography indicative of archaic introgression as the Neanderthal is more closely related to the derived haplotypes and the Denisovan to the ancestral haplotypes (Figure 3). Instead this indicates that the common ancestor of the three hominins already had polymorphisms formed in the *FADS* region, allowing for greater differentiation between modern humans as well as in the archaics. When the major haplotypes are displayed in a neighbor joining tree by genetic distance, this hypothesis is further supported (Figure 4). The Neanderthal has the ancestral haplotype, however it also has the overall haplotype topography that is more representative of the modern human derived haplotype than the modern human ancestral haplotype (Figure 3, 4, Additional File 7: Figure S5, Additional File 8: Figure S6). This indicates that the haplotype became polymorphic in the common ancestor of these three hominins.

**Figure 4.**
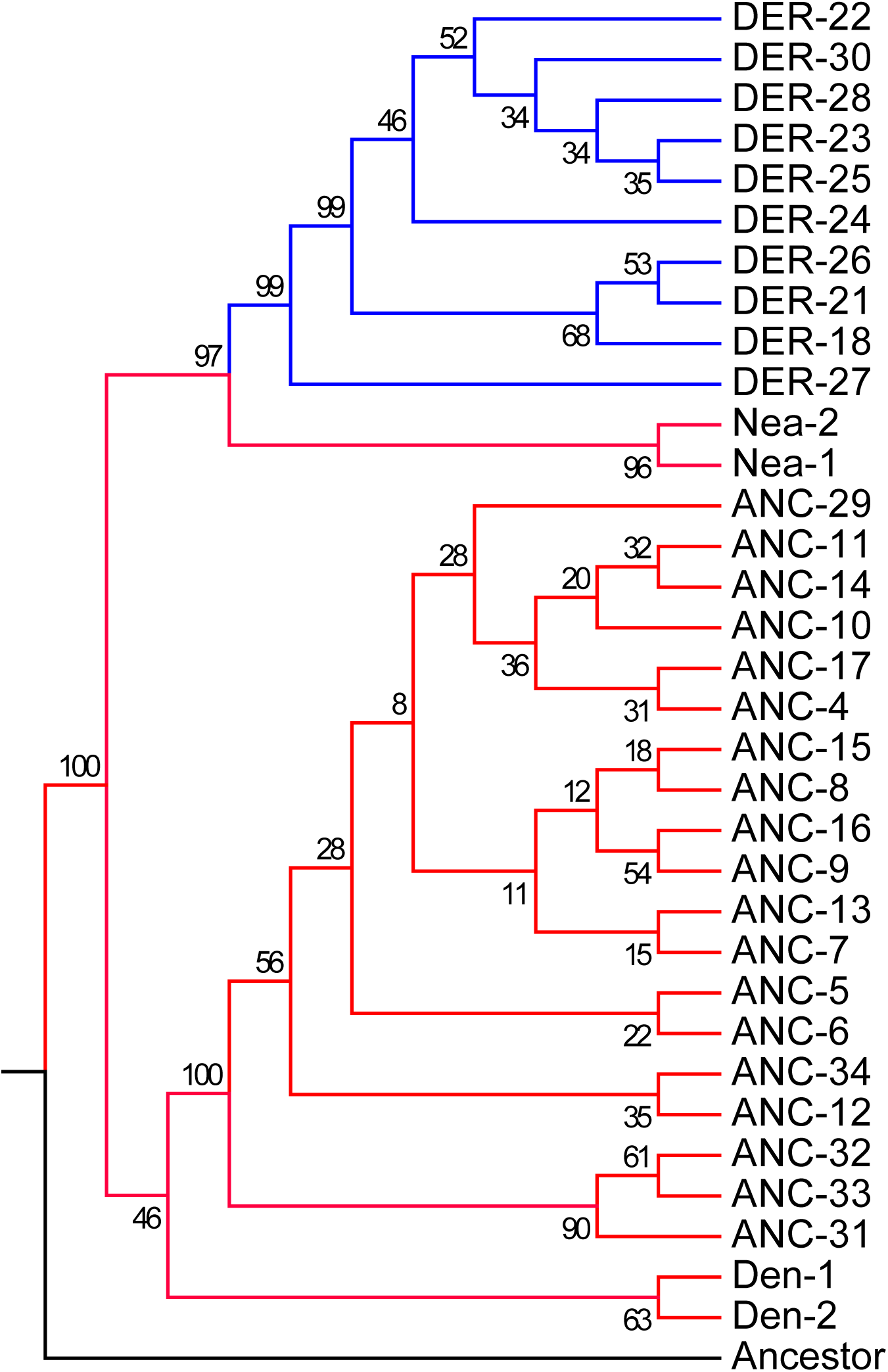
Tree representation of major haplotype networks, including invariant sites. The numbers on each node represent the percent that node was formed out of 500 bootstraps. Haplotype number is abbreviated as Hap# with the Number coming from the Roman numeral in the high coverage haplotype network (Additional File 8: Figure S6). Nea-1 = Haplotype 2, Nea-2 = Haplotype 3, Den-1 = Haplotype 19, Den-2 = Haplotype 20, and Ancestor = Haplotype 1

### 6. Derived Haplotype TMRCA

The derived haplotypes have a TMRCA of 688,474 ya (95% CI = 635,978 – 743,052). This is older than prior estimates by Ameur et al. [16] that showed the upper limit of the derived haplotypes to be only 433,000 ya. The inferred TMRCA value is within the estimated range of modern and archaic hominin divergence of 550,000 to 765,000 ya [22]. This estimate is robust to different values of human-chimpanzee divergence estimates (Additional File 9: Figure S7). Nearly all human-chimpanzee divergence values result in a TMRCA that is within the divergence of the three hominins, except for a human-chimpanzee divergence ≤ 5,192,642 which further supports that the derived haplotype began to differentiate within the modern human-archaic divergence. This would explain the great differentiation between the archaic and modern humans in this *FADS* haplotype because this would allow nearly 600,000 to 700,000 years of mutation to accumulate in this region on both the ancestral and derived haplotype. This also explains the Neanderthal haplotype being more closely related to the derived than the ancestral haplotypes as the differentiation began during the divergence of Neanderthals and modern humans.

## Discussion

Our results support the hypothesis that the different haplotypes in the *FADS* gene cluster represent an old polymorphism (Figure 3, Figure 4). We found no evidence to support the alternative explanation that the ancestral haplotype was re-introduced to non-Africans due to archaic introgression from Neanderthals or Denisovans. The TMRCA of the derived haplotypes is within the modern-archaic human divergence [22], however the haplotype topography and tree (Figure 3, 4) indicate that the TMRCA should predate the divergence of these hominins. Since both the ancestral and derived haplotypes are under positive selection (Figure 2) [15, 17-19], we are likely underestimating the TMRCA of the derived haplotypes. Positive selection causes a reduction in diversity of variants within a haplotype [71] and therefore a reduced TMRCA estimate. As a result, it is possible the actual TMRCA of the derived haplotypes does pre-date the modern-archaic human divergence. However the TMRCA we present is older than prior estimates [15, 16]. The differences in TMRCA calculations are likely due to each study using a slightly different genomic segment of the *FADS* region [15, 16]. In addition, we used a greater number of samples to calculate the TMRCA than Ameur et al. [16], which will likely increase haplotype diversity and lead to an older TMRCA. Further, we only used high coverage sequences (Unpublished Data^a^) [13] which allows us to identify more variants and therefore lead to a greater estimate of TMRCA than prior studies [15, 16]. Due to the old TMRCA of the derived haplotype, it is clear these *FADS* polymorphic haplotypes represent an old region of the hominin genome. This can be seen today by the great differentiation within and between the Neanderthal, Denisovan, and modern human ancestral and derived haplotypes (Table 1, Figure 3, and Figure 4). The Neanderthal haplotype clusters closer to the derived haplotype but do not share all variants as that of the core derived haplotype (Figure 3, Figure 4), which raises the question of whether the Neanderthal had full capability to synthesize LC-PUFAs from 18C-PUFAs. The old age of the derived haplotype further [13] establishes the importance of the *FADS* gene cluster in hominin evolution because it began to form within the divergence of modern humans and Neanderthals and Denisovans [22].

While prior work was done on a few Native American ancestry samples [13, 15, 16], with 386 Native American haplotypes, we determine that the Native American populations are near fixed for the ancestral haplotype. This is puzzling due to the potentially detrimental health effects that could arise from having a reduced capability to synthesize LC-PUFAs that are vital to brain and immune system development [1, 2]. However, Fumagalli et al. [17] suggest that the ancestral haplotype could be linked to cold weather adaptation possibly as a response to dietary restrictions of a cold climate. We further support this selection pressure due to finding Siberian populations are at a high proportion for the ancestral haplotype and that the proportion increases the further North a population lives in Asia (Additional File 5: Figure S4). According to one recent hypothesis, Native American ancestors remained isolated (possibly in Beringia) for up to 10,000 years prior to migrating into North America [72, 73], where there would have been a strong selection pressure for genetic variants that assisted in adapting to cold weather. Therefore, if this haplotype aids in cold weather adaptation, then Native American ancestors’ journey through Siberia and Beringia would favor the ancestral haplotype being pushed to high frequency, which is further supported by Amorim et al. [18]. In addition, some have posited that Native American ancestors spread into North America through a coastal route [74]. This could have facilitated reduced selection for LC-PUFA biosynthesis due to the fact that large quantities of LC-PUFA would have been found in seafood along the coast [8]. Eventually Native American populations left the coast and moved inland, however the persistence of the ancestral haplotype could be explained by the low effective population size of Native American populations (Unpublished Data^a^). Their low effective population size would require an extremely high selection pressure to reduce the haplotype frequency in a population already with the haplotype at near fixation [75]. To further evaluate this,more research is required on Native American populations that live far from major water sources and away from a cold weather climate, such as the Pima [76].

We were unable to observe the same two large LD blocks as seen by Ameur et al. [16]. This is likely due to us observing little to no recombination over the entire *FADS* gene region (Additional File 1: Figure S1). In addition, Ameur et al. [16], used a larger dataset of European ancestry individuals (5,652 individuals) which likely substantially increased their power to detect LD blocks compared to the 114 Native American ancestry individuals from the Peruvian Genome Project in our study. It is also possible that the LD block is specific to European ancestry as LD block formation are not as strongly seen in African ancestry cohorts [14]. However we were able to identify one small block in the Peruvian samples, which indicates that if we were to increase our sample size, that we could see the previously identified LD blocks [16].

LC-PUFAs are essential for a wide range of human biological functions [3-6]. A reduced capacity to synthesize LC-PUFAs has the potential to be a public health risk for modern populations with high Native American ancestry. For example, the n-3 LC-PUFA, DHA is known to be critical for brain function throughout the human life span, but its accumulation is especially important to healthy brain development during gestation and infancy [2, 4]. In the brain, DHA has a wide range of neurological functions including membrane integrity, neurotransmission, neurogenesis and synaptic plasticity, membrane receptor function and signal transduction [2]. Additionally, n-3 LC-PUFAs such as DHA, DPA and EPA and their metabolites have potent anti-inflammatory properties [1, 2]. There has been a dramatic increase in dietary exposure to the n-6 18C-PUFA, LA due to the addition of vegetable oil products to the MWD over the past 50 years [7]. This increase has shifted the ratio of n-6 to n-3 18C-PUFAs ingested to greater than 10:1 which assures that n-6 LA and not n-3 ALA is the primary substrate that enters the LC-PUFA biosynthetic pathway thereby producing ARA and not EPA, DPA and DHA [7]. So the critical question from a gene-diet interaction perspective is; does the near fixation of the ancestral haplotype with its limited capacity to synthesize LC-PUFAs in Native American ancestry individuals together with an overwhelming exposure of LA relative to ALA entering the biosynthetic pathway give rise to n-3 LC-PUFA deficiencies and resulting diseases/disorders in Native American populations? Simply stated, what are the sources of n-3 LC-PUFAs for Native Americans during critical periods of brain development and as anti-inflammatory mediators [77]? Questions such as these indicate that future research is needed to assess circulating and tissue total PUFA levels in Native American ancestry individuals. If these individuals are found to have low LC-PUFA levels, then they will be an important cohort to study the risk of LC-PUFA deficiencies and related dietary interventions in this area could provide a substantive benefit to Native American ancestry populations’ medical care.

## Conclusions

Through the analysis of Native American ancestry individuals from the Peruvian Genome Project (Unpublished Data^a^), all 1000 Genomes Project individuals [30], and African and European American individuals from GeneSTAR [13], we can confidently state that Native American local ancestry haplotypes are nearly fixed for the ancestral haplotype. Further, we fail to support the hypothesis that the ancestral haplotype was re-introduced to non-African populations through archaic introgression. Instead, we support the hypothesis that this *FADS* haplotype is an ancient polymorphism that first became polymorphic during the divergence of modern humans from Neanderthals and Denisovans with likely different selection pressures maintaining the variation in different parts of the globe. In addition, the near fixation of the ancestral haplotype in Native American ancestry populations together with shifts in 18C-PUFA exposure in the MWD raises the likelihood that *FADS* gene-dietary PUFA interactions could lead to numerous diseases/disorders and thus demands future efforts to analyze this potential public health problem impacting Native American ancestry populations.

### Abbreviations

ALA =: α–linolenic acid
ARA =: arachidonic acid
C =: carbon
CI =: confidence interval
DHA =: docosahexaenoic acid
DPA =: docosapentaenoic acid
EPA =: eicosapentaenoic acid
FADS =: fatty acid desaturase
LA =: linoleic acid
LC =: long chain
MWD =: modern western diet
n3 =: omega-3
n6 =: omega-6
PUFA =: polyunsaturated fatty acid
TMRCA =: time to the most recent common ancestor
ya =: years ago

## Declarations

### Ethics approval and consent to participate

Not applicable

### Consent for publication

Not applicable

### Availability of data and materials

Peruvian Genome Project variant calls are available upon request to be made to T.D.O.

## Author Contributions

T.D.O., R.A.M., F.H.C. conceived of the project. D.N.H performed all bioinformatics analyses. R.A.M., I.R. L.R.Y performed data generation and quality control analyses on GeneSTAR samples. R.A.M., L.B., D.B. were responsible for the recruitment of the GeneSTAR samples. All authors contributed to the writing of the paper.

## Acknowledgements

We thank Mait Metspalu, director of the Estonian Biocentre, Tartu, Estonia for providing geographic coordinates for the Siberian populations. We also thank Joana C. Silva, Amol Shetty and Michael Kessler for helpful discussion about results and experimental design.

## Endnotes

a. Daniel N. Harris, Wei Song, Amol C. Shetty, Kelly Levano, Omar Cáceres, Carlos Padilla, Víctor Borda, David Tarazona, Omar Trujillo, Cesar Sanchez, Michael D. Kessler, Marco Galarza, Silvia Capristano, Harrison Montejo, Pedro O. Flores-Villanueva, Eduardo Tarazona-Santos, Timothy D. O’ Connor, Heinner Guio. The Evolutionary Genomic Dynamics of Peruvians Before, During, and After the Inca Empire.

